# Volatile-mediated signaling induces resistance of barley against infection with the biotrophic fungus *Blumeria graminis* f.sp. *hordei*

**DOI:** 10.1101/2021.12.08.471267

**Authors:** Silvana Laupheimer, Reinhard Proels, Sybille B. Unsicker, Ralph Hückelhoven

## Abstract

Plants have evolved a vast variety of secondary metabolites to counteract biotic stress. Volatile organic compounds (VOCs) are carbon-based molecules induced by herbivore attack or pathogen infection. A mixture of plant VOCs is released for direct or indirect plant defense, plant-plant or plant-insect communication. Recent studies suggest that VOCs can also induce biotic stress resistance in distant organs and neighboring plants. Among other VOCs, green leaf volatiles (GLVs) are quickly released by plant tissue after the onset of herbivory or wounding.

We analysed VOCs emitted by 13-day old barley plants (*Hordeum vulgare* L.) after mechanical wounding using passive absorbers and TD-GC/MS detection. We investigated the influence of pure (Z)-3-hexenyl acetate (Z3HAC) as well as complex VOCs from wounded barley plants on the barley - powdery mildew interaction by pre-exposure in a static and a dynamic headspace connected to a powdery mildew susceptibility assay.

GLVs dominated the volatile profile of wounded barley plants with Z3HAC as the most prominent compound. Pre-exposure with Z3HAC resulted in induced resistance of barley against fungal infection. Barley complex volatiles emitted after mechanical wounding, similarly, enhanced resistance in receiver plants.

We found volatile-induced modification of the interaction towards an enhanced resistance against fungal infection. In addition, Z3HAC triggered a modulation of the alcohol dehydrogenase isoenzyme activity in receiver plants, a physiological response that possibly contributes to induced resistance. Plant-originated volatile metabolites could be a useful supplementation for future agronomic or horticultural practices.

**Highlight:** Volatile-induced modification of the barley-powdery mildew interaction towards an enhanced resistance against fungal infection.

## Introduction

Volatile organic compounds (VOCs) are known as signals in plant communication (Ninkovic *et al*., 2021). They attract pollinators or enhance plant defense against insect herbivores and pathogens. The emission of VOCs is either constitutive or stress-induced. In the latter case, amount and composition of VOCs depends on the kind of damage (Ameye *et al*., 2018; Mithöfer *et al*., 2009; Paré & Tumlinson, 1996). According to Dudareva *et al*. (2005) VOC response caused by biotic stresses has various functions: inter- and intraspecific plant communication, allelopathy or as cross kingdom chemical signals for herbivores or for natural enemies of the attacking herbivores. In addition, volatile emission by herbivore-damaged plants may not only directly trigger defense responses, but instead prime the plant itself and neighbouring plants for higher responsiveness upon further herbivore attack (Baldwin & Schultz, 1983; Ton *et al*., 2007; Ye *et al*., 2021). A close connection between phytohormones and VOC biosynthesis is given in plants (Scala *et al*., 2013). Jasmonic acid (JA) and its derivatives are related to the class of green leaf volatiles (GLVs), both synthesised *via* the octadecanoid pathway. Phytohormones might play a crucial role in plant-pathogen interaction because a burst of JA and other stress-related defense hormones and genes could be observed after herbivory or infection by fungal pathogens (Ameye *et al*., 2015; Kessler & Baldwin, 2002).

GLVs are ubiquitously produced by all green plants (Hatanaka, 1993) and constitutively emitted in small amounts from intact and unstressed plant tissue (Piesik *et al*., 2011). Nevertheless, predominant synthesis and release of GLVs occurs after tissue damage, like herbivory or wounding. GLVs might act as wound signals in plants and also as intra- and interspecific messengers (Bate & Rothstein, 1998; Shiojiri *et al*., 2006).

Engelberth *et al*. (2004) revealed the ability of plant priming with three pure synthetic GLVs, (*Z*)*-*3-hexenal, (*Z*)*-*3-hexenol, and (*Z*)*-*3-hexenyl acetate. GLV-treated corn (*Zea mays cv. Delprim*) induced higher JA concentrations in response to herbivory than control plants. A similar response was shown in wheat (*Triticum aestivum*) pre-treated with pure (*Z*)*-*3-hexenyl acetate (Z3HAC) and challenged with the hemibiotrophic fungus *Fusarium graminearum* (Ameye *et al*., 2015).

Most studies investigated the role of GLVs on plant-insect interaction and studies on the potential of GLVs in plant-pathogen interaction focused on necrotrophic fungi (Finiti *et al*., 2014; Kravchuk *et al*., 2011; Scala *et al*., 2013; Vicedo *et al*., 2009). The biotrophic powdery mildew is one of the most common fungal pathogens of cereals (IS Compendium, 2020). However, little research exists to investigate resistance modulation by naturally occurring external signals, like GLVs, of cereal crops against biotrophic fungi.

Here, we use the biotrophic ascomycete *Blumeria graminis* f. sp. *hordei* (*Bgh*) and its natural host barley (*Hordeum vulgare* L.) (Wyand & Brown, 2003) as a model system. The epiphytic stages of *Bgh* are well known and allow an easy monitoring of the fungal development on plant tissue (KIta *et al*., 1981; Schulze-Lefert & Vogel, 2000). Fungal spores grow with a defined program upon leaf inoculation. After penetration of the host cell wall, *Bgh* develops a specialized hyphal structure to enter the intact epidermal cell and create a cellular interface within 24 hours. This so-called haustorium is mainly for nutrient uptake from alive host cells and for effector delivery (Panstruga, 2003), and it likely is the only structure that serves to feed the pathogen. Hence, intact host leaf tissue is required. After this successful fungal establishment, a formation and further growth of mycelium on leaf tissue occurs 3-7 days after infection (Eichmann & Hückelhoven, 2008). Diverse forms of induced resistance are known for the barley pathosystem, and respective gene expression or physiological markers have been described (Beßer *et al*., 2000; Dey *et al*., 2014; Käsbauer *et al*., 2018; Kogel & Langen, 2005). The main intrinsic function of plant alcohol dehydrogenases (ADHs) is in anaerobic fermentation during hypoxia, but they also play a critical role in the biosynthetic pathway of GLVs (Strommer, 2011). In GLV biosynthesis, C_6_- and C_9_-aldehydes, alcohols, and esters originate from lipid degradation in the chloroplast. Membrane-bound hydroperoxide lyases catalyse the release of two structurally different aldehydes into the cytosol, namely hexanal and (*Z*)-3-hexenal. An isomerase factor is responsible for the spontaneous conversion of (*Z*)-3-hexenal to its isomer (*E*)-2-hexenal (Scala *et al*., 2013). The next step is the reduction of the aldehyde to its corresponding alcohol by ADHs. The alcohols are further processed by alcohol acyltransferases that catalyse the formation to the corresponding acetate ester (Matsui, 2006). GLVs vary in their toxicity and volatility. Aldehydes are most toxic and least volatile. They exert insecticidal as well as fungicidal effects (Kishimoto *et al*., 2008; Nakamura & Hatanaka, 2002). Barley has three fermentative *ADH* genes (Harberd & Edwards, 1983; Trick *et al*., 1988): The *HvADH1* gene is constitutively expressed in seeds, shoots, and roots under aerobic conditions, whereas *HvADH2* and *HvADH3* are only expressed under stress or when oxygen is insufficient (Good *et al*., 1989). ADH enzyme activity is regulated at the transcriptional level and can be induced by abiotic factors, like oxygen deficiency, drought, or salt stress (Kennedy *et al*., 1992; Myint *et al*., 2015; Xiong & Schumaker, 2002) and biotic factors. Proels *et al*. (2011) reported a critical function of the ADH-dependent fermentative metabolism in plant-pathogen interactions. Infection of barley *cv*. Ingrid with *Bgh* results in an induction of ADH enzyme activity and also *HvADH1* and *HvADH2* gene expression is enhanced. Additionally, RNAi-mediated transient knock-down of *HvADH1* revealed a reduction in the fungal penetration success (Pathuri *et al*., 2011). Systemic resistance in barley induced by a chitin elicitor was associated with reduced ADH activity and stable over-expression of *HvADH1* in barley inhibited chitin-induced resistance. ADHs hence may play a critical role in the response to biotic stresses and can potentially serve as a physiological marker for induced resistance responses in barley (Käsbauer *et al*., 2018).

Here, we show that a pre-exposure of young barley *cv*. Golden Promise or Ingrid with different concentrations of Z3HAC leads to enhanced resistance to subsequent challenge-inoculation with powdery mildew spores. The VOC profile emitted after mechanical wounding caused similar induced resistance in the receiving plant. A potential role of ADH in the physiological response to VOCs is discussed.

## Methods

### Plant material

Experiments were performed with 13-day old barley plants (*Hordeum vulgare* L. *cv*. Golden Promise and Ingrid). Seeds were germinated for 3 days in the dark and then transferred to non-sterilised soil (CL ED73, Werke Patzer, D). Depending on the type of experiment, 10 - 15 individuals were grown together in a 7 × 7 cm plastic pot and watered every day. Plants were cultivated in the climate chamber with a 16:8 (light:dark) photoperiod at a temperature of 18 ± 2°C, a light intensity of 150 ± 2 µE*s^-1*^m^-2^ and a relative humidity of 65 %.

### Cultivation of *Blumeria graminis* f.sp. *hordei*

The powdery mildew fungus *Blumeria graminis* (DC.) Speer f. sp. *hordei* (*Bgh*) Em. Marchal race A6 served as pathogen. It was maintained on barley *cv*. Golden Promise in a climate chamber MLR-351H (Sanyo, Moriguchi, JPN) at a temperature of 18 ± 2°C, a light intensity of 150 µE*s^-1*^m^-2^ and 65 % of relative humidity.

### VOC collection

Volatiles in the static headspace of barley plants were collected with polydimethylsiloxane (PDMS) tubes. The PDMS tubes were prepared as described in Kallenbach *et al*. (2014). Briefly, PDMS hoses (1 mm internal diameter x 1.8 mm external diameter, PDMS, Carl Roth) were cut into 5 mm long pieces and stored in 4:1 (v/v) acetonitrile/methanol. After 3 h incubation at 25°C the PDMS pieces were transferred out of the solvent into a 200 ml glass column containing a glass frit. The column was connected to a nitrogen gas flow with 5 l/min to flush solvent and contaminants and then heated under the nitrogen flow for 1.5 h at 210°C. After cooling the PDMS tubes were transferred into 4 ml screw-neck glass vials containing argon. The vials were screwed tightly and sealed with polytetrafluorethylene (PTFE). Three PDMS tubes were bead on an acetone cleaned Ø 0.05 mm aluminium wire and hung into the oven bag without touching the bags. VOC collections were performed under constant conditions in the climate chamber ATC26 (Conviron, Winnipeg, CAN) for 24 h. Qualitative analysis of volatiles emitted by barley *cv*. Golden Promise were done 1, 3, 5.5, 8, 10.5, 13 and 24 h after wounding. Thereby, the first and the second leaf of 8 to 10 individuals was heavily damaged with a bluntish razor blade applying consecutive cuts and crushes every few millimeters across the entire leaf blade. Additionally, VOCs were collected in the same way from non-wounded control plants and pots without any plant as a system control. Immediately after wounding or inoculation, an oven bag with a size of 50 × 31 cm (Toppits, Minden, DE) was wrapped around each of the barley plants (including the pot) and tightly closed with a cable binder at the upper end. An elastic band was installed right at the upper rim of the pots. After volatile collection, the PDMS tubes were transferred to 1.5 ml screw neck glass vials and stored at -20°C until further analysis.

### VOC analysis and identification

VOCs trapped with PDMS tubes were detected by gas chromatography coupled to a quadrupole mass spectrometer (GC-MS-QP2010 Ultra, Shimadzu, Kyoto, JPN). At first, PDMS tubes were placed in 89 mm glass TD tubes (Sigma Aldrich, St. Louis, USA) and analytes were released from the adsorbent by an ultra thermo desorption unit (TD20, Shimadzu). After desorption in He with a flow rate of 60 ml/min at 200°C for 8 min, the substances were cryo-focused onto a trap (Tenax®-GR adsorbent resin, Tenax, Baltimore, USA) at -20 °C.

This trap was heated to 230°C within 10 sec and analytes were injected on a GC column with a length of 30 m, a diameter of 0.25 mm and a film thickness of 0.25 µm (Rtx-5MS, Restek, Bellefonte, USA). As carrier gas helium was used with a constant linear velocity of 44.3 cm/s. The GC program was set to 45°C for 3 min followed by 180°C with a thermal increase of 6°C/min. At least the temperature was increased to 320°C with 100°C/min and was held for 15 min to clear the column.

Mass spectrometry data of the samples were detected with the method of electron impact within the spectrum from 33 to 350 m/z at 70 eV and a scan speed of 1666 Da/s. MS data were analysed and named with the program GCMS solution, Version 4.20 (Shimadzu, 2014) based on the Wiley Registry of Mass Spectral Data, 8th Edition (Shimadzu, 2006) and NIST11 database (National Institute of Standards and Technology, Gaithersburg, Maryland, USA). The adjustment of each compound and a unique method file with a compound list containing target ions (m/z) at a specific retention time were created by hand. The peak area was integrated automatically by the program using this method file. Results were evaluated in Integrated Peak Area per g fresh weight (IPA/g FW).

### VOC-mediated signaling in a static or dynamic headspace

A receiver test with a single VOC were performed in two connected hard plastic tubes with a total volume of 2.2 litres. This setup allows a defined headspace and an indirect and consistent treatment of the whole plant with the volatile compound of interest. Pure standards of (*Z*)*-*3-hexenyl acetate (Z3HAC, 98 %, CAS number 3681-71-8, Merck) was used in different concentrations. After VOC application on a filter paper in one of the plastic tubes, both tubes were connected *via* a pipe and tightly closed and sealed (supplemental Figure S1). The experiment ran in the climate chamber MLR-351H (Sanyo, Moriguchi, JPN) with a temperature of 18°C, a light intensity of 150 ± 2 µE*s^-1*^m^-2^ and a relative humidity of 65 %. 10 barley individuals per pot were exposed to 0.1 µM or 10 µM Z3HAC for 24 hours. Three individuals were pooled for protein studies. Six individuals were harvested and used for *Bgh* susceptibility assays. The remaining individual was disposed.

A similar setup was performed with a dynamic headspace sampling system. Two oven bags were connected by a pump system, which allows an active VOC treatment under controlled conditions (Tholl *et al*., 2006). Activated charcoal filtered air was continuously pushed into the first bag with 700 ml/min and an airflow of 400 ml/min was constantly pulled out of the first bag and pushed into the second bag, which contains the receiver plant. The second bag was not tightly closed to avoid accumulation of air or humidity. A higher influx flowrate was necessary to avoid air contaminations from the outside. The first bag contains either emitter plants (15 individuals per pot) or Z3HAC (0.1 µM or 10 µM) carried on a filter paper. The total volume of this system was 30 litres (supplemental Figure S2).

### *Blumeria graminis* f.sp. *hordei* susceptibility assay

14-day old barley plants exposed to Z3HAC, or the bouquet of wounded emitter plants were used in a susceptibility assay against *Bgh*. After exposure, either the leaves were cut and transferred to a 0.5% (w/v) agar plate with the adaxial side facing up or the intact individuals remaining in the pot were inoculated. To ensure a homogeneous inoculation with *Bgh* spores, leaves were completely spread out and placed flat on the agar by additionally fixing them with small metal rods (Ø 2 mm). Inoculation took place in an inoculation tower (BxTxH 50×50×200 cm) with Ø 6-10 spores/mm². After 3 days incubation in the climate chamber ATC26 (Conviron, Winnipeg, CAN) the leaf segments were evaluated by macroscopically counting fungal microcolonies developing into *Bgh* pustules. For bleaching, leaves were transferred in a 15 ml Falcon tube containing 3:1 ethanol:acetic acid. After 3 days of incubation at room temperature and sunlight pustules could be coloured with ink (Hückelhoven & Kogel, 1998; Linkmeyer *et al*., 2013). Results were calculated in developing pustules/cm^2^.

### Quantification of ADH isoenzyme activity

After exposure with different concentrations of Z3HAC in a static headspace, 2^nd^ leaves were collected as pooled samples of 3 individuals. Plant tissue was harvested, frozen in liquid nitrogen and ground using the tissuelyzer II (Qiagen). After crude protein extraction samples were centrifugated twice at 12500 rpm for 10 min at 4°C. Supernatant was transferred in a 1.5 ml reaction tube and kept on ice.

After Bradford quantification (Bradford, 1976) samples mixed with loading dye were loaded on the polyacrylamide matrix (Mini-PROTEAN® Tetra System 12-0625 BioRad, Hercules, California, USA). To detect the separated ADH isoforms, an activity staining was performed using Nicotinamide adenine dinucleotide (NAD^+^), ethanol and N-nitroblue tetrazolium (NBT). For evaluation of equal loading of proteins and normalistion for further analysis, the gels were stained with Coomassie RAPIDStain™ (Biosciences). The Fusion SL Darkroom Concept connected to the DarQ7 camera (Vilber Lourmat, Marne-la-Vallée, FR) were used to capture images of the native gels and perform a relative quantification of stained ADH isoenzymes (Proels *et al*., 2011). Image acquisition, analysis and quantification were performed semi-automatically with the program FusionCapt Advance SL4, Version 16.03 (Vilber Lourmat, 2011). Results were normalized to the total protein band, which was stained with Coomassie, and evaluated in percent to the control. Results are given in ADH activity in % to control.

### Data analysis

All data were checked for statistical assumptions, i.e. homogeneity of variances and normal distribution. The VOC data were very variable and therefore, the random forest algorithm was investigated (Breiman, 2001). Analysis was performed with RStudio, Version 0.98.978 (RStudio, Inc. 2013). For data of developing *Bgh* pustules and ADH isoenzyme activity Welch’s correction unpaired t-test was conducted using GraphPad Prism® 6 (Version 6.01).

## Results

### (*Z*)-3*-*hexenyl acetate dominates the VOC bouquet of mechanically wounded barley

To evaluate the dynamic behavior of VOC emission in 13-day old barley plants, untreated plants were compared to mechanically wounded plants. VOCs were collected with passive absorbers over a time frame of 1 to 24 hours after wounding in a closed headspace. This method allows for a qualitative identification of the compounds and a semi-quantitative assessment of emission levels based on *Integrated Peak Area normalized on gram fresh weight* (IPA/g FW).

Generally, young barley plants emitted only a few VOCs. Only 11 compounds, originating from the barley plant, could be detected in untreated and wounded plants (supplemental Table S1, supplemental Figure S3). Thereby, the GLVs were highly induced after mechanical wounding and dominated the volatile blend. Beside terpenoids barley plants constitutively emitted a certain amount of the following GLVs: (Z)-3-hexenal, (Z)-3-hexenol (Z3HOL), (E)-2-hexenol and (*Z*)-3*-*hexenyl acetate (Z3HAC). Directly after mechanical wounding the amount of total emitted GLVs increased around 10-fold (33600 IPA/g FW) compared to the untreated control plants (3600 IPA/g FW). We recorded the maximum of GLV emission at 3 hours post wounding (hpw) with 52000 IPA/g FW. There was a significant increase in VOC emission following mechanical wounding.

To get a closer look on individual GLVs, the emission over time of the most dominant GLV, Z3HAC, was analysed. Z3HAC is the end product of a typical GLV biosynthetic pathway and the most volatile compound (Hatanaka, 1993; Maffei, 2010). Z3HAC was significantly induced immediately after mechanical wounding with 25000 IPA/g FW compared to the non-treated control. The maximum was reached after 3 hpw with 40000 IPA/g FW (Figure 1). Subsequently, we recorded a successive decrease of the amount of Z3HAC until 24 hpw. An analysis of the whole volatile blend at 1 hpw classified Z3HOL as the most important compound discriminating between wounded and control plants by the random forest algorithm (supplemental Figure S3). Beside the GLVs, 2,3-butanediol, butane and the terpenoids, β-caryophyllene and α-terpinolen, could be detected. Both terpenoids also showed a dynamic emission from 1 to 3 hpw with a lower descriptive pattern than the dominant GLVs. After 3 h Z3HAC accounted for the highest fraction of the complex volatile bouquet (supplemental Figure S3).

**Figure 1:**
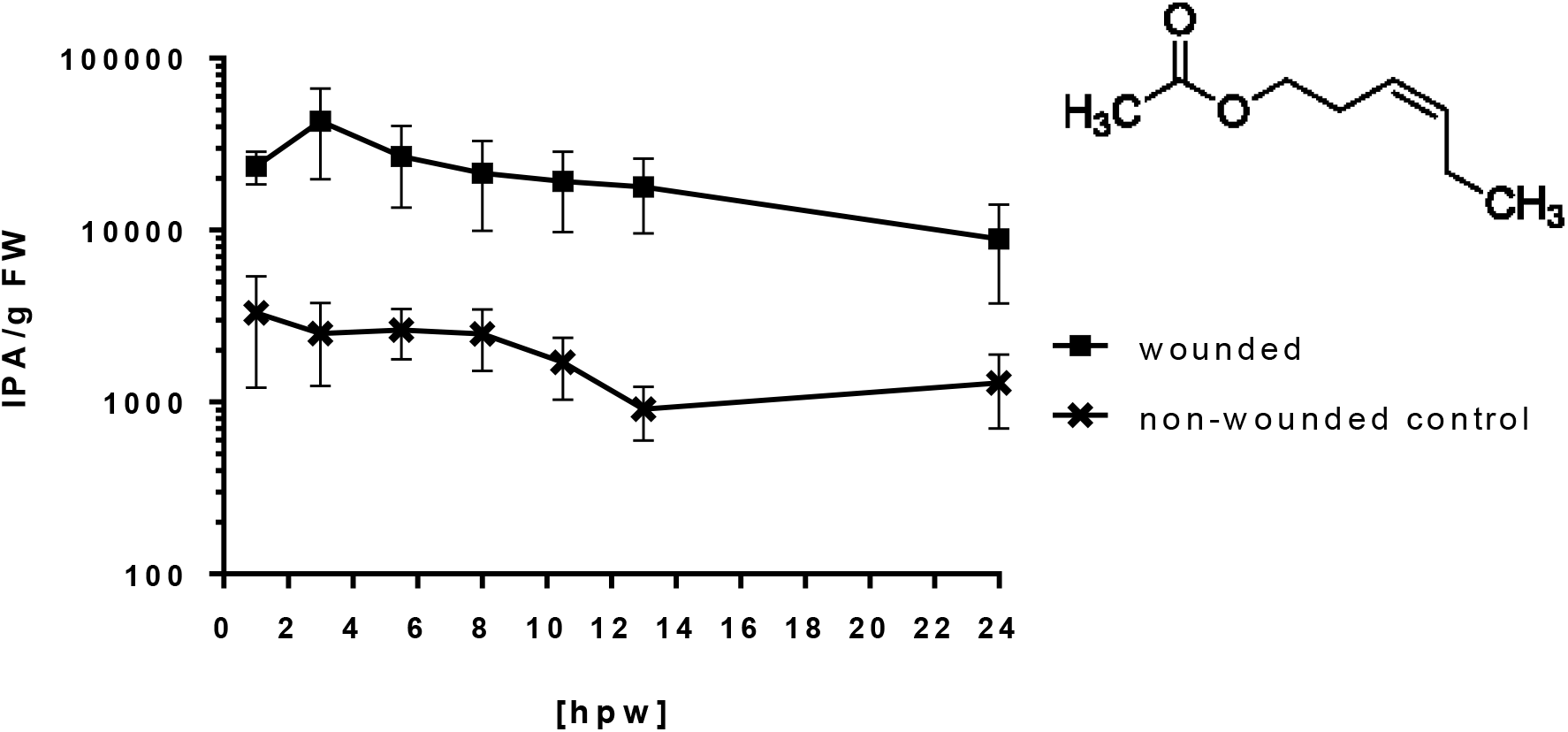
Chemical structure and emission rate of (Z)-3-hexenyl acetate (Z3HAC) after mechanically wounding in comparison to a non-wounded control over time from 1 to 24 hours post wounding (hpw). VOCs were collected in a closed headspace with passive absorbers and identified via TDU-GC/MS (n = 7). For wounding, the first and second leaf were strongly damaged with a blunted razor blade, thereby applying consecutive cuts and crushes every few millimeters alongside the leaf blade. Y-axis shows Integrated Peak Area (IPA) per g fresh weight (g FW). Error bars represent means ± SE.

### VOC profiles modulate barley resistance against *Bgh*

The most prominent compound in the volatile composition of 13-day old barley was Z3HAC. Hence, we used pure synthetic Z3HAC in a VOC exposure experiment to test if disease resistance of young barley plants could be modified with one particular VOC. Thereby, we focused on the second leaf of 13-day old barley plants as well as exposure times of 24 or 48 hours.

Barley plants (*cv*. Golden Promise) were placed in a static headspace and exposed for 24 h to 0.1 µM or 10 µM of synthetic Z3HAC. The compound was applied in one step on a filter paper within the closed headspace, defining the start point of the experiment. The concentration of 0.1 µM was calculated by reference to previous quantitative volatile collection experiments in barley and may resemble the emission rate, when young barley plants are mechanically wounded (supplemental Figure S4).

Five individuals of the receiver plants were used for *Bgh* inoculation on excised leaves. 14-day old barley leaves were sampled after 24 h exposed to Z3HAC and prepared for inoculation on agar plates. Fungal microcolonies with strongly branched epicuticular hyphae developing into pustules were counted 3 days after inoculation with *Bgh*. This revealed a significantly reduced number of developing pustules and consequently reduced susceptibility of barley to powdery mildew with both applied concentrations (Figure 2).

**Figure 2:**
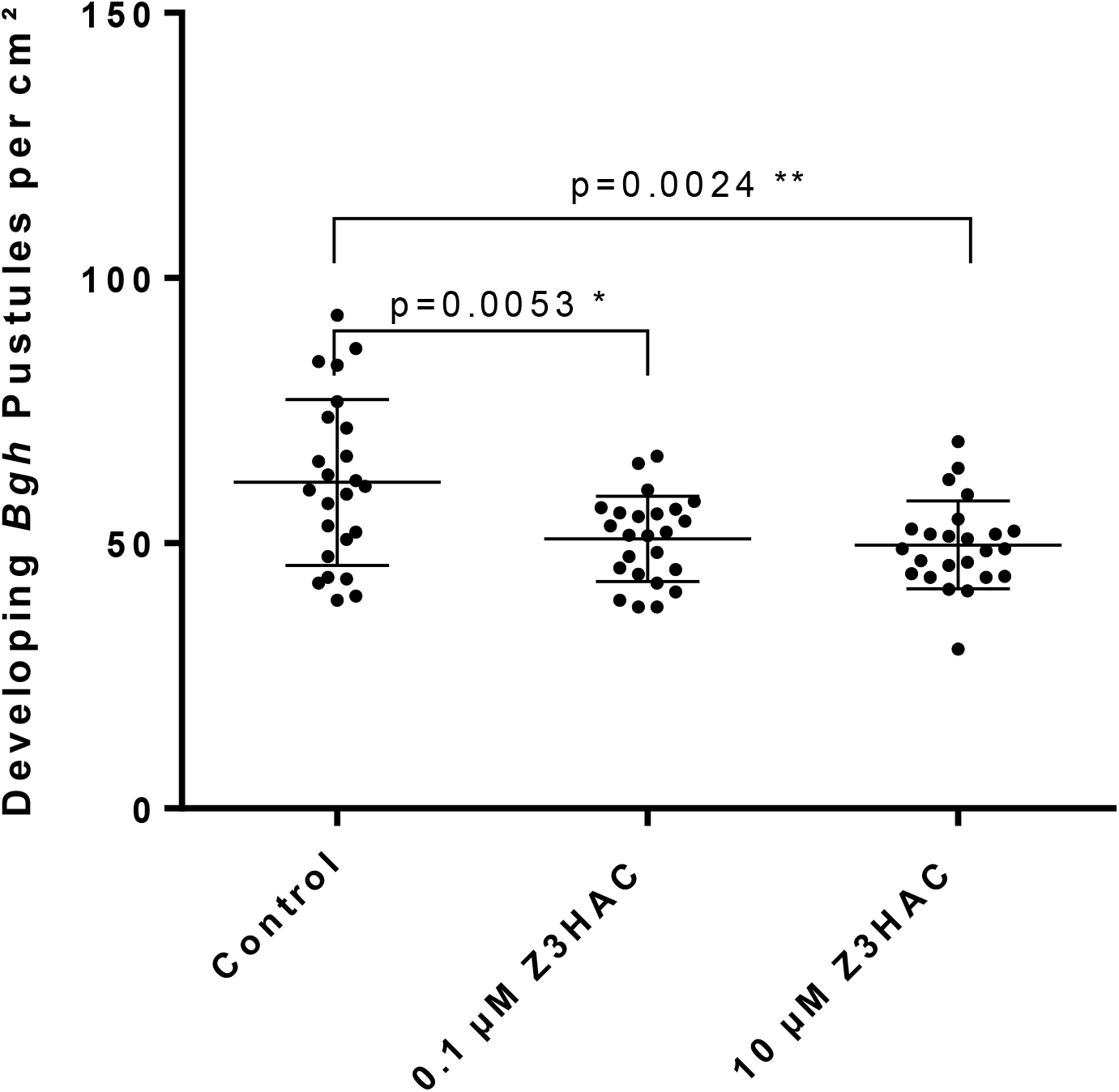
Developing Bgh Pustules on 2^nd^ leaf of young barley plants (cv. Golden Promise). 13-day old barley plants were exposed to 0.1 µM or 10 µM Z3HAC for 24 h in a static headspace. Afterwards 2^nd^ leaves were collected and inoculated with *Bgh* spores on agar plates. 72 h after inoculation, microcolonies developing *Bgh* pustules were counted (n=24). Data show developing *Bgh* pustules per cm^2^ leaf area compared to a non-pre-exposed control. Error bars represent means ± SD (p < 0.005 ** or 0.05 *; Welch’s unpaired t-test).

To avoid physiological side-effects of a closed headspace lacking gas exchange with the environment, such as increasing humidity or CO_2_ depletion, we further used a dynamic headspace sampling system with a defined flux of in- and out coming air. Pots with 15 plants of barley *cv*. Ingrid were exposed to 0.1 or 1 µM Z3HAC for 48 h. First approaches with 1 µM Z3HAC in a static headspace revealed a significant reduction of pre-treated plants compared to control plants (supplemental Figure S5). By using a dynamic and open system, the concentration of Z3HAC is expected to decrease rapidly after application. Hence, a dilution factor must be considered. With a constant flow rate of 700 ml/min influx and 400 ml/min outflow it is theoretically expected that the concentration within the headspace will drop to 0.01 µM within 30 min. Therefore, we applied a higher concentration of 1 µM Z3HAC in the same dynamic setup for 48 h exposure time. In this setup, both dosages of Z3HAC again induced resistance in receiver plants against powdery mildew. The observed effects were very similar in strength when compared to the static system. We then adapted the *Bgh* susceptibility assay to avoid further effects of cutting the leaves off receiver plants. The individual leaves remained attached to intact plants in the pot for the inoculation and incubation time of 72 h. Only then, leaves were excised to count the number of established fungal microcolonies. The cultivar Ingrid responded in a similar way as seen for Golden Promise by induced resistance to powdery mildew when pre-treated with Z3HAC. The 2^nd^ leaf showed a similar trend after 48 h of 0.1 and 1 µM Z3HAC exposure compared to non-treated control plants (Figure 3).

**Figure 3:**
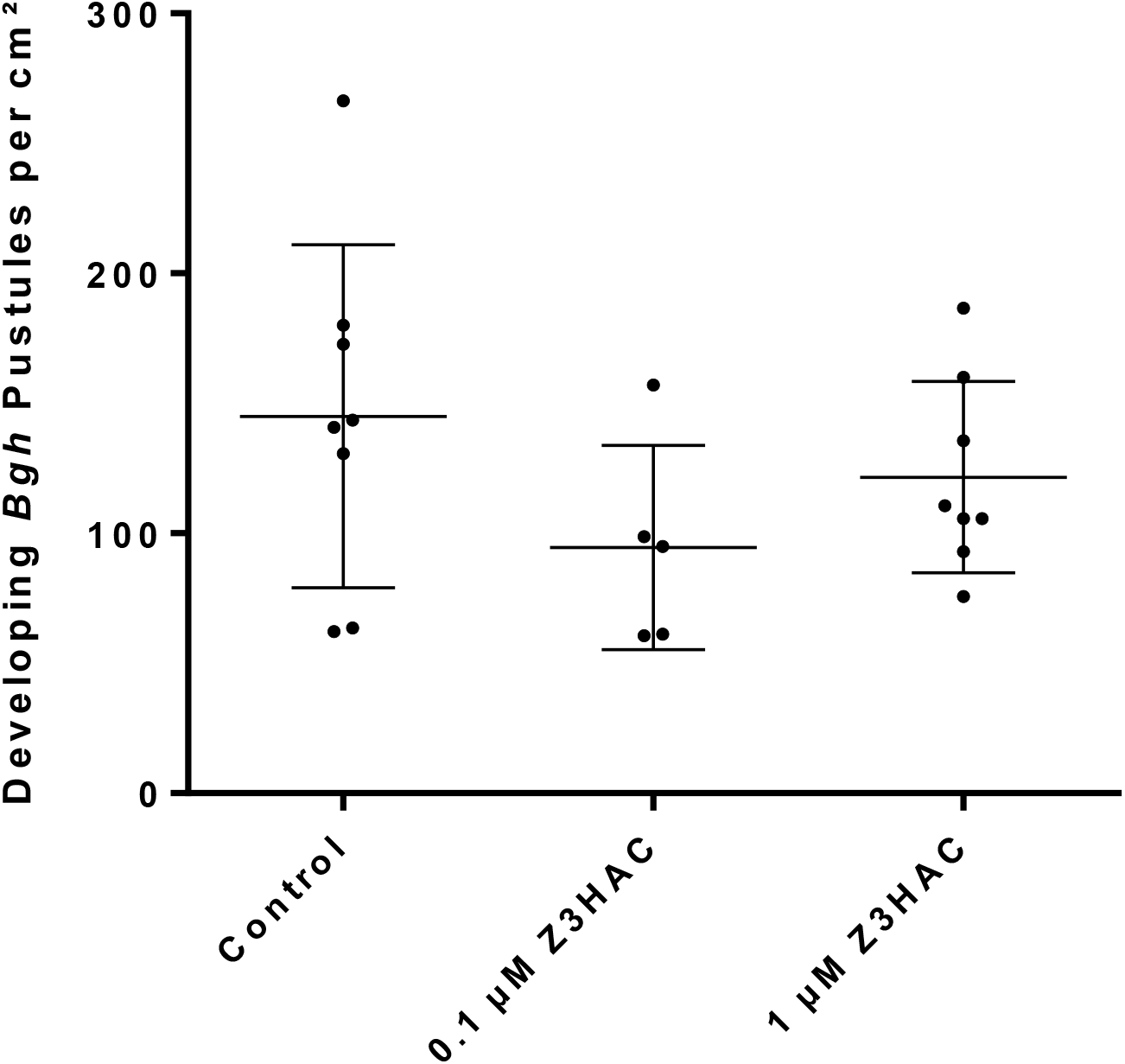
Developing Bgh Pustules on 2^nd^ leaf of young barley plants (cv. Ingrid). After 48 h exposure with 0.1 or 1 µM Z3HAC in a dynamic headspace the individuals remained intact in the pot and were inoculated with *Bgh* spores. 72 h after inoculation microcolonies were counted. Shown are developing *Bgh* pustules per cm^2^ leaf area compared to a non-pre-exposed control. Error bars represent mean ± SD with n=5 (0.1 µM) and n=8 (Control, 1 µM).

We were also interested if the complex bouquet of a wounded VOC-emitting plant could lead to a defense response in the receiving plants as seen for Z3HAC. Pots with 15 plants of *cv*. Ingrid were used as emitter and receiver plants. The experiment was performed in the dynamic headspace system for 48 hours. The emitter plants were mechanically damaged over the entire leaf blades, which marks the starting point of the exposure time. As a control, a non-wounded plant was used as emitter. After 48 h of pre-exposure, receiver plants were released from the headspace, remained attached to the intact plant in the pot and were inoculated with *Bgh* spores. 72 h after inoculation we observed a significantly reduced number of established microcolonies developing into powdery mildew pustules (Figure 4). This suggests a modification of the defense response of receiver plants, when pre-exposed to the complex volatile profile of mechanically wounded barley plants.

**Figure 4:**
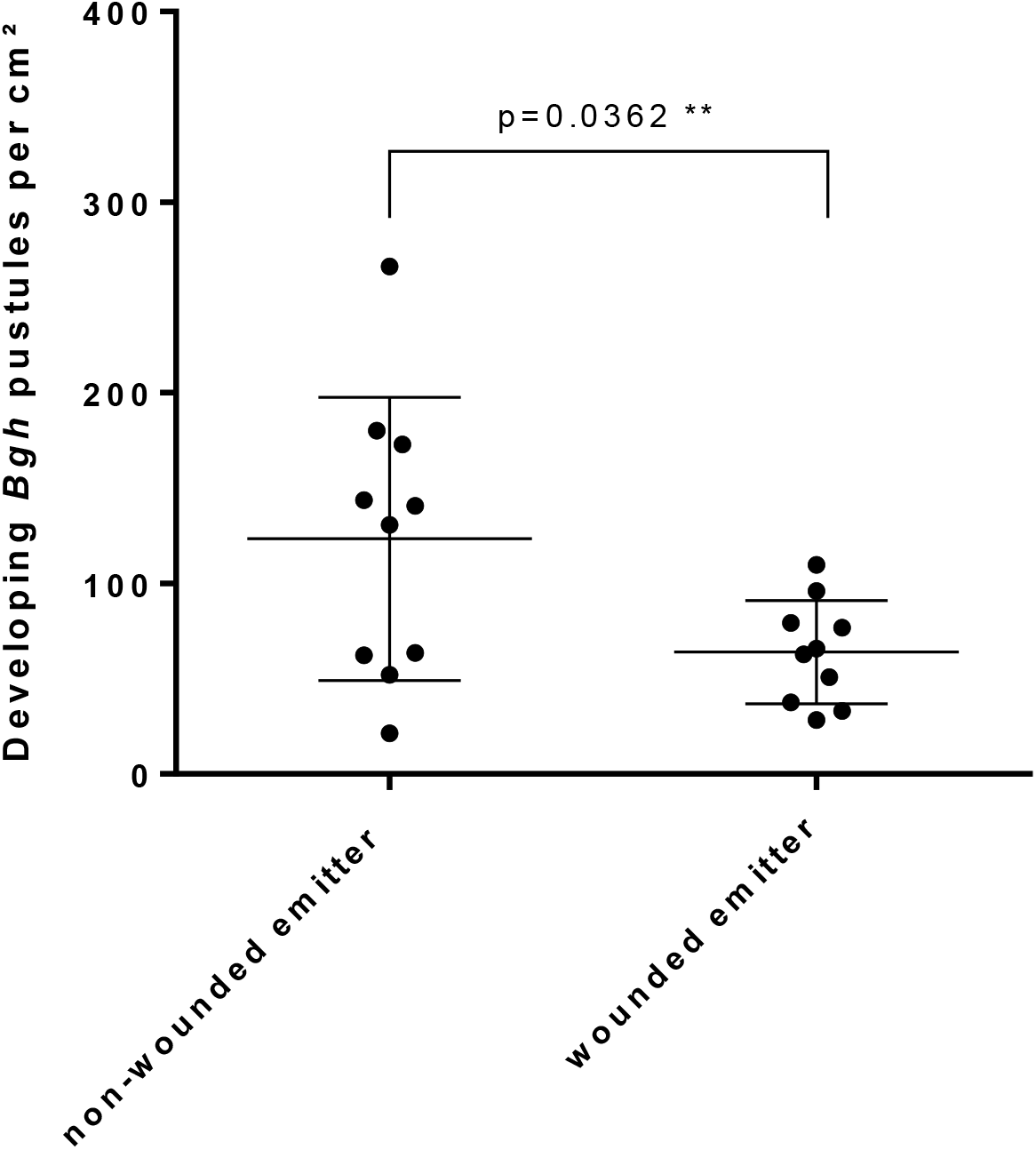
Developing Bgh Pustules on 2^nd^ leaf of barley plants (cv. Ingrid). After 48 h exposure with the complex bouquet of wounded barley plants the individuals remained intact in the pot and were inoculated with *Bgh* spores. 72 h after inoculation, microcolonies were counted (n=10). Shown are developing *Bgh* pustules per cm^2^ leaf area compared to a non-wounded emitter control. Error bars represent mean ± SD (p < 0.05; Welch’s unpaired t-test).

### VOC profiles modulate ADH isoenzyme activity

To investigate the effect of Z3HAC on the ADH isoenzyme activity in barley, pre-exposed plants were used for a specific in gel ADH isoenzyme activity staining assay (Proels *et al*., 2011). All three active protein dimers (ADH 1-1, ADH 1-2 and ADH 2-2) showed a similar pattern of regulation when normalized to total protein bands in the gel poststained after ADH measurement. Intensity of *in gel* activity staining was evaluated and data are shown as percent (%) of the non-pre-exposed control, which was set to 100% (Figure 5a-c). Interestingly, a significantly reduced ADH enzyme activity was observed in the 2^nd^ leaf after exposure to 0.1 µM Z3HAC for the isoenzyme dimers ADH1-1 and ADH1-2, whereby ADH1-2 showed a strong effect with about 50 % residual activity. For 10 µM Z3HAC the effect of reduced ADH1-1 and ADH1-2 isoenzyme activity is weaker than the response to 0.1 µM Z3HAC. When considering only 0.1 µM Z3HAC as the concentration that likely represents physiological conditions best, the 2^nd^ fully grown leaf showed a consistent down-regulation of all ADH isoform activities. The dimer ADH2-2 showed for both concentrations a similar tendency of reduced enzyme activity in the 2^nd^ leaf.

**Figure 5:**
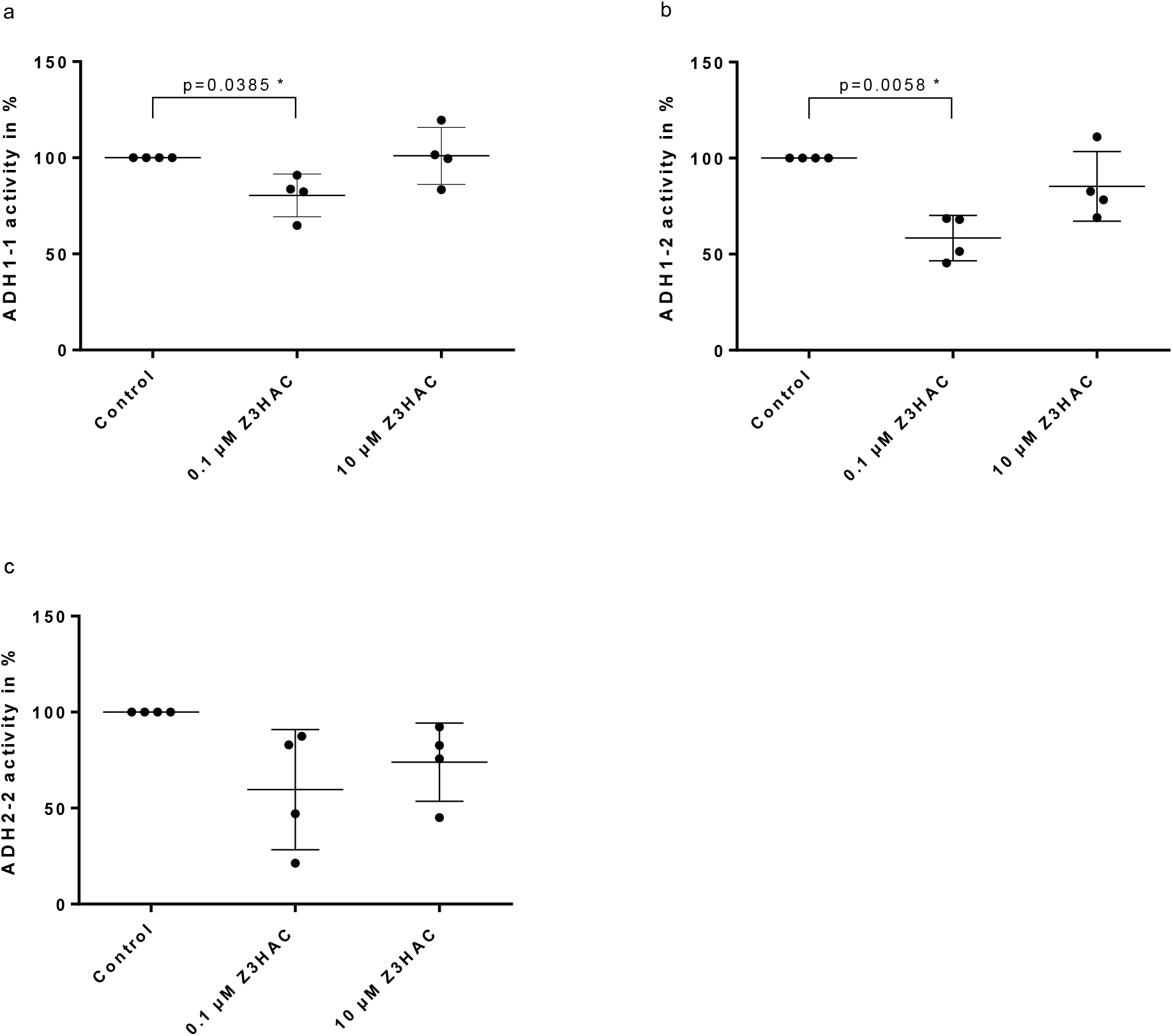
ADH isoenzyme (ADH1-1 (a), ADH1-2 (b) and ADH2-2 (c) activities in the 2^nd^ leaf. 13-day old barley *cv*. Golden Promise were placed in a static headspace and passively exposed with 0.1 µM and 10 µM Z3HAC. After 24 h leaf material was collected as pooled samples of 3 individuals. Proteins were extracted and ADH isoenzyme activity were determined (n=4). Shown are ADH activity in percent (%) of the non-pre-exposed control (100%). Error bar represents mean ± SD (p < 0.05; Welch’s unpaired t-test).

We repeated this experiment in a dynamic headspace system. In addition, we prolonged the exposure time to 48 hours to see if the enzyme activity is changed compared to 24 hours exposure. For 48h of exposure to Z3HAC, all three isoenzymes showed the same pattern of downregulation in the 2^nd^ leaf (Figure 6a). Enzyme activity was significantly reduced for the ADH1-2 dimer and the mean over all three isoenzyme dimers. The results after exposure to the VOC bouquet of a wounded plant differed. Here, the three ADH isoenzyme dimer activities varied in the 2^nd^ leaf and were similar or increased when compared to controls (Figure 6b). Together, these findings suggest a role of the single compound Z3HAC as an external signal that provokes a physiological response in barley and induces resistance to *Bgh*. ADH activity sensitively reacts to leaf exposure to Z3HAC. Low to moderate concentrations of Z3HAC suppress leaf ADH activity in most experimental setups.

**Figure 6:**
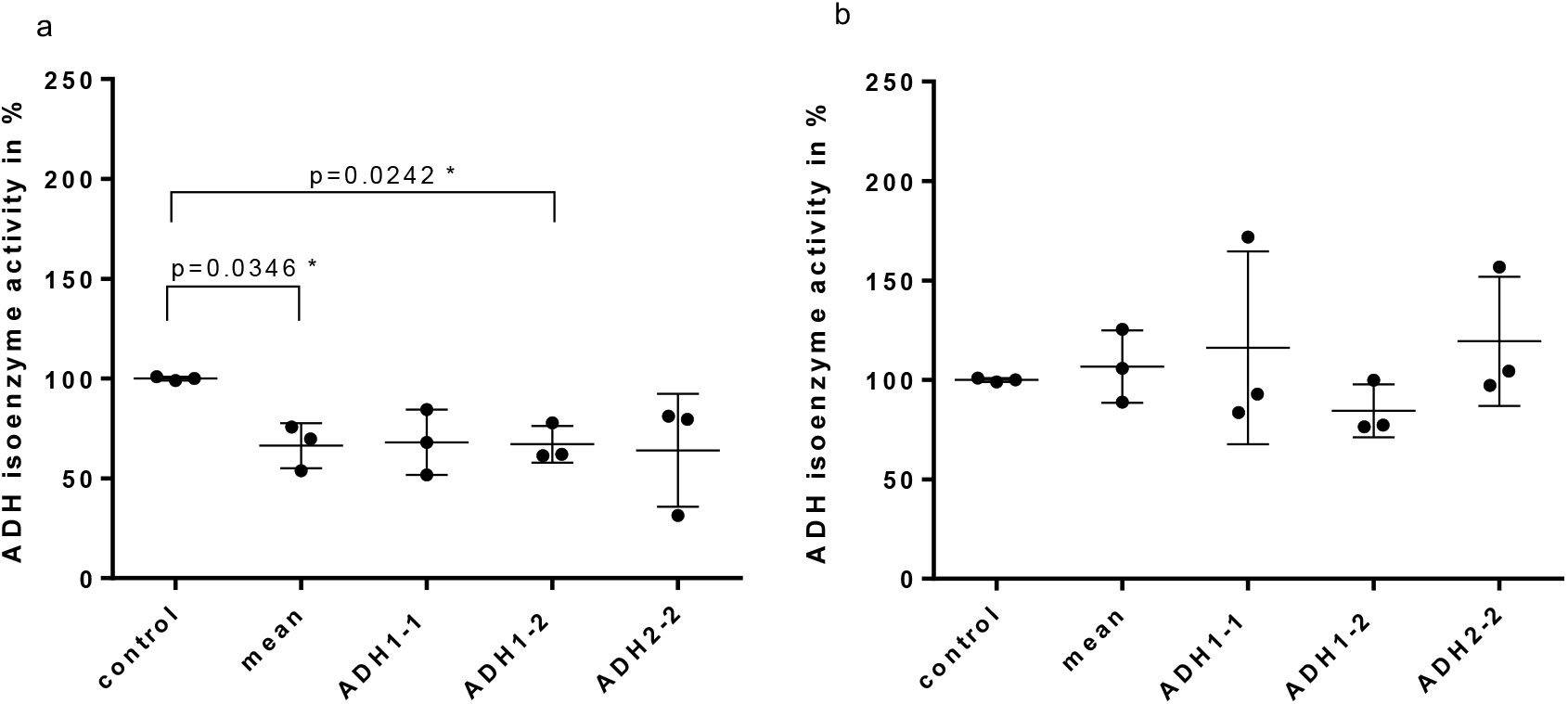
ADH isoenzyme activities in the 2^nd^ leaf after different types of pre-exposure. 13-day old barley *cv*. Ingrid were placed in a dynamic headspace and exposed to either 1 µM Z3HAC (a) or the complex bouquet of wounded emitter plants (b). After 48 h leaf material was collected as pooled samples of 5 individuals. Proteins were extracted and ADH isoenzyme activity were determined (n=3). Dots show individual ADH isoenzyme activity in percent (%) of the non-pre-exposed control (100%). Mean shows the average activity over all three isoenzymes. Error bars represent mean ± SD (p < 0.05; Welch’s unpaired t-test).

## Discussion

The volatile profile of 13-day old barley plants *cv*. Golden Promise was identified, and mechanically damaged plants mostly emit GLVs. These VOC group are known as common chemical defense measures of plants and therefore, they are focused on in research for biocontrol of pests and diseases. Here we found that exposure to the synthetic GLV Z3HAC or the bouquet of a wounded emitter plant enhances the resistance of young barley plants to infection with *Bgh*. Pre-treated plants showed a reduced susceptibility of barley to *Bgh* infection compared to non-treated control plants and two efficient Z3HAC dosages were observed. Pre-exposure with Z3HAC also led to a modulated ADH isoenzyme activity, a potential physiological marker for induced resistance to *Bgh*.

The composition of VOCs released by young barley plants in different stress situations are hardly known. Most studies focussed on later physiological stages and grains of cereals or root tissue to characterise VOCs emission (Delory *et al*., 2016; Gfeller *et al*., 2013; Piesik *et al*., 2011). Here we investigated the volatile blend of 13-day old barley plants *cv*. Golden Promise in untreated and mechanically damaged plants. GLVs dominated the total volatile blend with Z3HAC as a typical example of this VOC class. Piesik *et al*. (2010) observed a similar composition after mechanical wounding in barley *cv*. Rasik. However, the amount and ratio of emitted compounds differed. This might be caused by the different cultivars and the developmental stages. We elucidated a highly dynamic emission of GLVs within a time frame of 24 h. In the first 3 h after mechanical damage the maximum was reached, and until 24 h the emission of Z3HAC gradually decreased. These observations are comparable to those from earlier studies. Apparently, the mode and frequency of damage influences GLV emission (Ameye *et al*., 2015; Mithöfer *et al*., 2009), and acetates are the most volatile compounds within the group of GLVs (Hatanaka, 1993; Maffei, 2010). GLVs are suggested as general responses after biotic and abiotic stresses and therefore might play an important role in plant responses to stressful environments. Signaling between plants or distinct plant organs has been reported to be mediated by volatiles. However, the biological function of GLVs from signal perception to signal transduction are still poorly understood. Nevertheless, they likely can act as signals to activate defense responses in surrounding plants.

Induced resistance in barley against *Bgh* has already been shown after abiotic stresses, like heat (Vallelian-Bindschedler *et al*., 1998) osmotic stress, or drought (Wiese *et al*., 2004), and also for chemical compounds, such as Chitosan and benzothiadiazole (Beßer *et al*., 2000; Faoro *et al*., 2008). An induction of resistance triggered by volatile organic compounds, especially GLVs, has been evidenced in maize (Engelberth *et al*., 2004), lima bean (Kost & Heil, 2006), poplar (Frost *et al*., 2008), tomato (Finiti *et al*., 2014) and wheat (Ameye *et al*., 2015). However, most studies investigated the role of GLVs against herbivore attack, while research on crops and their fungal diseases are rare. Despite the fact that barley is the fourth most-produced cereal in the world, little information is available on functions of GLVs in interaction with biotrophic leaf pathogens such as *Bgh*.

We exposed barley plants to a volatile set emitted by mechanically wounded plants or a single compound, Z3HAC, in different concentrations. After 24 or 48 hours, we challenged these plants with *Bgh* spores. First developing microcolonies, indicative of successful fungal development, were visible after three days. We observed in independently repeated experiments different responses due to the physiological age of the leaf material (Torres *et al*., 2017) and inconsistent results with exposure times of just 6 hours. By contrast, Z3HAC exposure for 24h resulted in reproducible outcomes. Pre-exposures in static headspaces decrease the fungal infection success on the second leaf. For 48 hours, the volatile profile of a mechanically wounded plant led to a significantly reduced number of 50 % developing pustules compared to a non-pre-exposed control plant. Exposure of 0.1 or 10 µM Z3HAC for 24 h significantly reduced the susceptibility of barley against *Bgh* infection. These observations could be repeated in a dynamic headspace with 0.1 and 1 µM Z3HAC for 48 h. Preliminary data suggest that higher concentrations of 100 µM were inefficient in inducing resistance. Thus, the VOC composition, exposure time, dose and the headspace volume may have an important impact on the resistance response.

GLVs are an important group of VOCs and produced in plants upon different types of stresses. They are involved in a range of interactions and can act as antibacterial and antifungal compounds (Kishimoto *et al*., 2008; Nakamura & Hatanaka, 2002). Also priming of plants by modulating phytohormonal levels and stress-related genes has been shown (Ameye *et al*., 2015; Engelberth *et al*., 2004; Frost *et al*., 2008). These observed general responses and our observations suggest an important role of GLVs in induced resistance in barley. Volatile treatments impede the pathogen success on barley, but the mechanisms behind remain elusive. Rather the number than the size of developing pustules was reduced in VOC-pre-treated barley. This may make a direct antifungal effect of VOCs unlikely. From earlier studies of this pathosystem, it can be deduced that this is rather explained by a reduced success in initial cell wall penetration and haustoria formation, which is the pivotal step in early pathogenesis (Eichmann and Hückelhoven, 2008). Further studies might address the exact mechanism of VOC-induced cellular or biochemical defence measures.

Previous investigation of barley and *Bgh* revealed a critical role of ADHs on the plant-pathogen interaction. *Bgh*-infection in barley *cv*. Ingrid led to an increased ADH activity in comparison to non-infected control. Further, a knock-down of *HvADH1* in transiently transformed barley epidermal cells or in stable transgenic plants leads to a reduced fungal penetration success, and *vice versa*, overexpression of *HvADH1* leads to an enhanced fungal penetration success. Additionally, the immunogenic elicitor chitin triggers systemic resistance in barley and systemic downregulation of the isoenzyme ADH1-1 activity. This chitin-induced resistance is limited when *HvADH1* is ectopically over-expressed (Käsbauer *et al*., 2018; Pathuri *et al*., 2011). We hence investigated whether ADH activity would reflect induced resistance of barley against *Bgh* and compared this in control and VOC-induced conditions.

We exposed plants for 48 hours with Z3HAC or the complex wounding bouquet. The treatment of 1 µM Z3HAC reduced the ADH activity of all three active isoenzyme dimers (ADH1-1, ADH1-2, ADH2-2), whereas the treatment of the complex GLV bouquet resulted in less consistent ADH activity compared to a non-pre-exposed control. Hence, ADH activity alone does not explain VOC-induced resistance in barley but indicates a physiological response of barley to VOCs. ADH activity by itself should contribute to the composition and amount of GLVs by converting aldehydes to alcohols, which are then substrates for alcohol acyltransferases that produce Z3HAC (Bate & Rothstein, 1998; Scala *et al*., 2013). It seems thus possible that ADH activity is part of a local and systemic rheostat that reacts to diverse kinds of GLVs in a possibly concentration-dependent manner. It is tempting to speculate that ADH thereby also contributes to GLV signaling by local adaptation of cellular physiology.

Up to date, it is unclear how plants perceive GLVs, and which subsequent signaling mechanisms VOCs activate. An application of a single compound may result in a different response compared to exposure to a complex volatile profile emitted by a wounded plant. The results seen in Figures 2 and 3 would support this assumption. The picture arises that particular VOCs or volatile profiles can induce resistance against microbial pathogens. In line with data from the literature, our results support a complex involvement of ADH activity in VOC production (Strommer 2011) and in VOC-modulated barley-powdery mildew interaction. For now, it remains unclear whether VOC-induced resistance to biotrophic fungal parasites is relevant in nature. However, data clearly show that VOCs provoke physiological responses in barley and modify susceptibility to powdery mildew. Hence, the spatial expansion of VOC perception in the neighbourhood of damaged barley tissue in a field remains to be shown but VOCs should be considered as potential resistance-inducing factors in barley. There is an increasing need for alternatives to chemical fungicide solutions for disease management. Plant-originated volatile metabolites could be exploited as a supplementation for future agronomic or horticultural glasshouse practices. This may need further research into underlying molecular mechanisms, controlled application technology and breeding for high VOC sensitivities or emission of efficient inducer bouquets.

## Abbreviations

GLV: Green leaf volatile
VOC: Volatile organic compounds
JA: Jasmonic acid
Z3HAC: (Z)-3-hexenyl acetate
Z3HOL: (Z)-3-hexenol
ADH: Alcohol dehydrogenase
*Bgh*: *Blumeria graminis* f.sp. *hordei*
EV: Empty vector
*HvADH*: Mutant of ADH isoenzymes in *Hordeum vulgare* L. background

## Acknowledgements

This research was funded by the Deutsche Bundesstiftung Umwelt (DBU).

## Supplement

**Figure S1:**
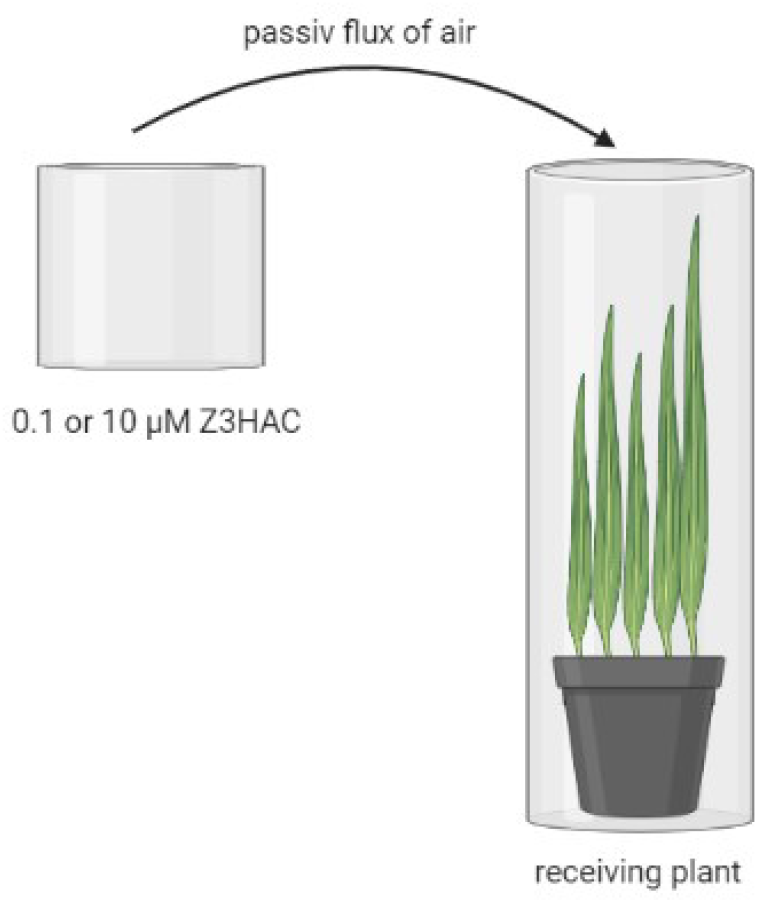
Schematic setup of VOC-mediated signaling in a static headspace. Two hard plastic tubes are connected by a pipe, which allows a passive flux of air. A filter paper carrying 0.1 or 10 µM Z3HAC is placed in the smaller tube as source of volatile emission. The bigger tube contains the receiving plant. Both tubes are tightly closed and sealed to avoid contamination from outside. The total volume of this system is 2.2 liters. Created with BioRender.com.

**Figure S2:**
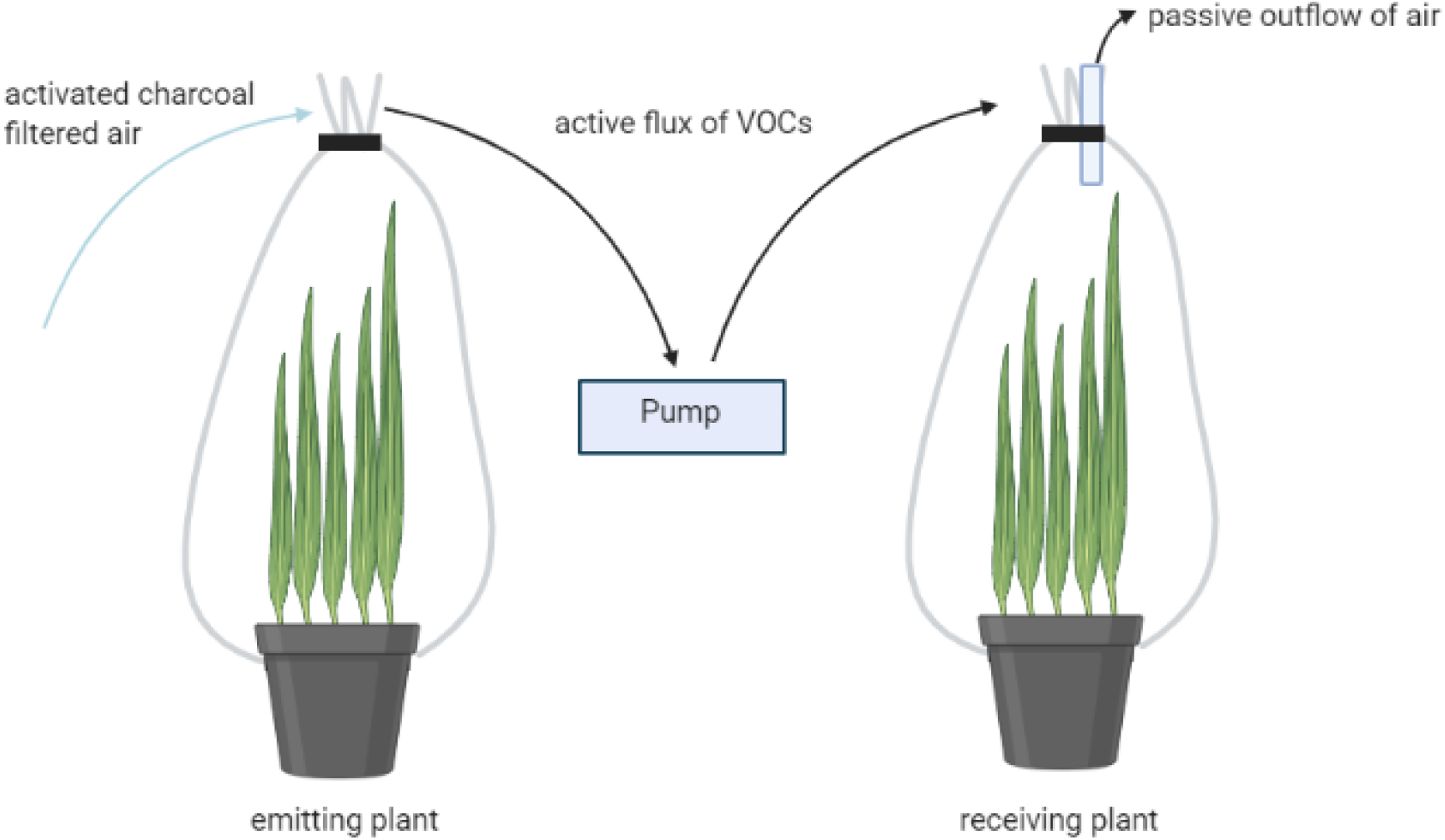
Schematic setup of VOC-mediated signaling in a dynamic headspace. Activated charcoal filtered air is actively pushed into the first bag containing the emitting plant, which is either a mechanically wounded plant or a non-wounded control plant. For pre-exposure with Z3HAC dosages the plant is replaced by a filter paper carrying 0.1 µM or 1 µM Z3HAC. The first bag is closed tightly at both sides including the pot and the soil. The influx into the first bag is higher than the outflow of air to avoid air contaminations from outside. Another pump pushes the air from the first bag into the second bag, which contains the receiving plant. The second bag allows a passive outflow of air to avoid accumulation of air and humidity. The total volume of this system is 30 liters. Created with BioRender.com.

**Table S1:** VOCs originated from untreated as well as wounded 13-day old barley plants cv. Golden Promise. Green leaf volatiles (GLVs), Terpenoids and Others (origin is unknown) were collected with passive absorbers (PDMS tubes) in a closed headspace and identified *via* TDU-GC/MS in untreated and wounded barley plants. For analysis, a unique method file for barley plants was established with the specific target ion (m/z) and the retention time (min), which are listed. Also the molecular weight in g/mol is given (Pubchem Open Chemistry Datebase, 2016).

| Class | Compound | Molecular Weight [g/mol] | Target Ion Mass [m/z] | Retention Time [min] |
| --- | --- | --- | --- | --- |
| GLVs | (Z)-3-hexenal | 98.14 | 69 | 4.547 |
|  | (Z)-3-hexenol | 100.16 | 67 | 5.825 |
|  | (E)-2-hexenol | 100.16 | 57 | 6.071 |
|  | Hexanoic acid | 116.16 | 60 | 8.553 |
|  | (Z)-3-hexenyl acetate | 142.2 | 82 | 9.682 |
|  | Heptanoic acid | 130.19 | 119 | 11.017 |
|  | Nonanoic acid | 158.24 | 83 | 16.370 |
| Terpenoids | α-terpinolen | 136.23 | 93 | 12.093 |
| | β-caryophyllene | 204.35 | 133 | 19.737 |
| Others | Butane | 58.12 | 58 | 2.052 |
|  | 2,3-butanediol | 90.12 | 57 | 4.415 |

**Figure S3:**
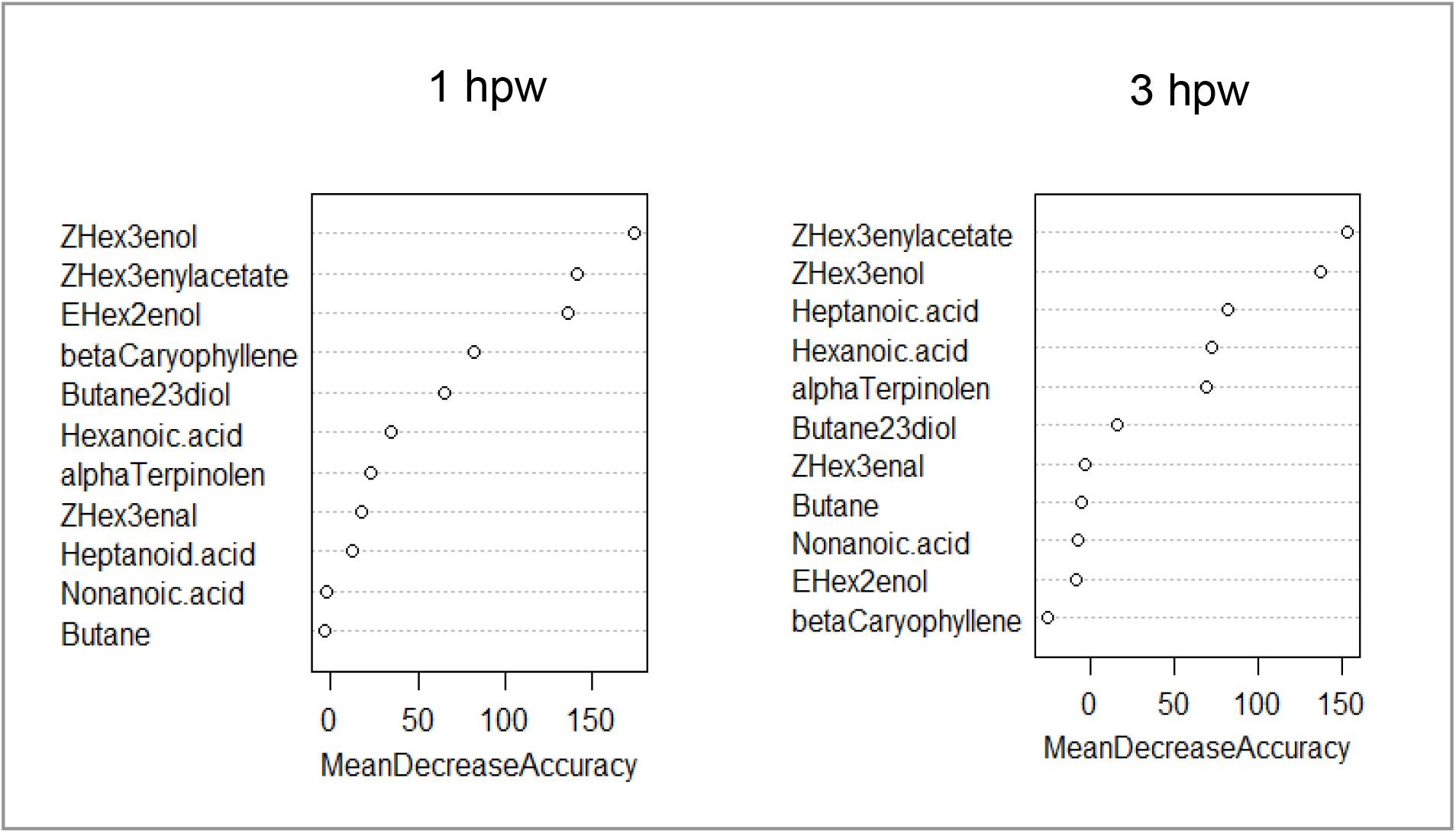
Relative importance of each VOC in the volatile blend of a 13-day old barley plants 1 or 3 hours after wounding (hpw) calculated by the random forest algorithm in R!. The listed compounds (Table S1) were emitted by the barley plant itself. A random forest algorithm sorted the 11 compounds in order to their relative importance in the complex volatile blend. Results are given in Mean Decrease Accuracy (MDA). VOCs were collected in a closed headspace with passive absorbers (PDMS tubes) and identified *via* TDU-GC/MS (n = 7).

**Figure S4:**
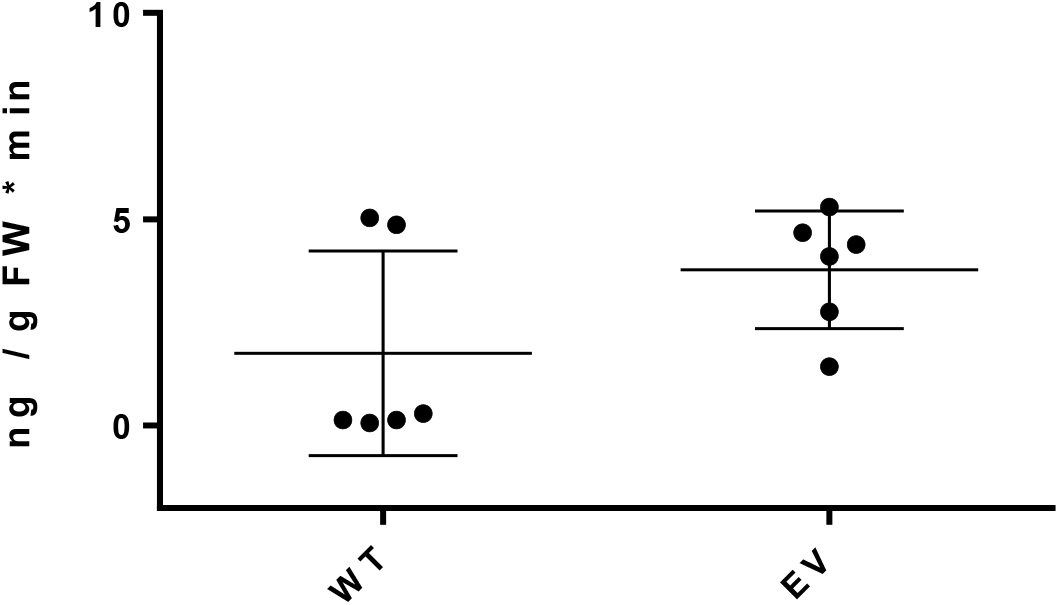
Emission rate of Z3HAC after mechanically wounding in Wildtype (WT) and empty vector (EV) as controls. To quantify the volatiles emitted from barley after wounding, a dynamic headspace sampling system installed in a climate chamber (York, Johnson Controls, Milwaukee, WI, USA) was used. This system allows a VOC collection under controlled conditions (Tholl *et al*., 2006). Activated charcoal filtered air was continuously pushed at 0.6 l/min into the bags and an airflow of 0.3 l/min was constantly pulled out of the bag over 120 min. The VOCs were trapped in a special glass filter containing 50 mg Super-Q absorbent (Analytical Research Systems, Gainesville, Florida, USA). After collection the absorbent was washed twice with 200 µl of dichloromethane (DCM) containing an internal standard (10 ng/µl Nonylacetate) in a 1.5 ml screw neck glass vial and stored at 4°C. Detection and analysis were performed by using two identically gas chromatographs (GC HP 6890, Agilent, Santa Klara, California, USA). One was connected to a mass spectrometry unit Agilent 5973 for detection of the fragmented molecular ions. The other was coupled to a flame ionisation detector (FID, H9200 hydrogen generator, Agilent) for quantification of the fragments. 1 µl of each analyt were injected on a GC column HP-5 (Agilent, 30 m length, 250 µm diameter, 0.25 µm film thickness) and the mobile phase was helium with a linear velocity of 51 cm/s. The initial temperature was 45°C for 2 min followed by 180°C with a thermal increase of 6°C/min. In the end the temperature was raised to 300°C for 2 min cleaning the column. MS data were detected with EI within the spectrum from 33 to 350 m/z at 70 eV and a scan speed of 1666 Da/s. The FID unit allows the quantification of the organic compounds using thermal ionisation of the molecules in a hydrogen flame (Otto, 2011). Chromatograms were visible with Agilent GCMS Enhanced ChemStation, Version E.02.00.493 (2008 Agilent) and comparison of these two chromatograms with the same retention times of one sample enables qualification and quantification of the compounds using the internal standards and the following equation. Error bars represent mean ± SD with n=6.

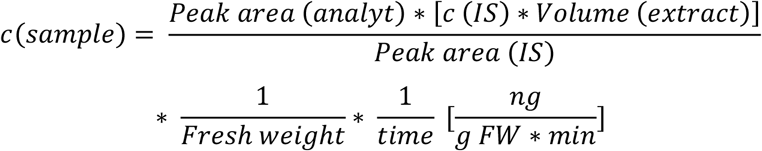 Calculated means are 1.8 ng/gFW*min for WT and 3.8 ng/gFW*min for EV. The approximated mean of 2.8 ng / g FW * min was used for calculating the expected concentration within the headspace with the following equation.

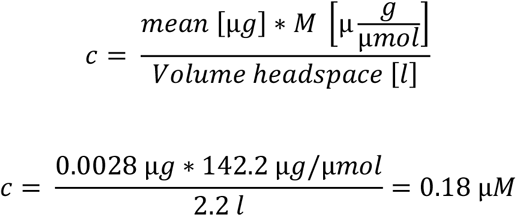

**Figure S5:**
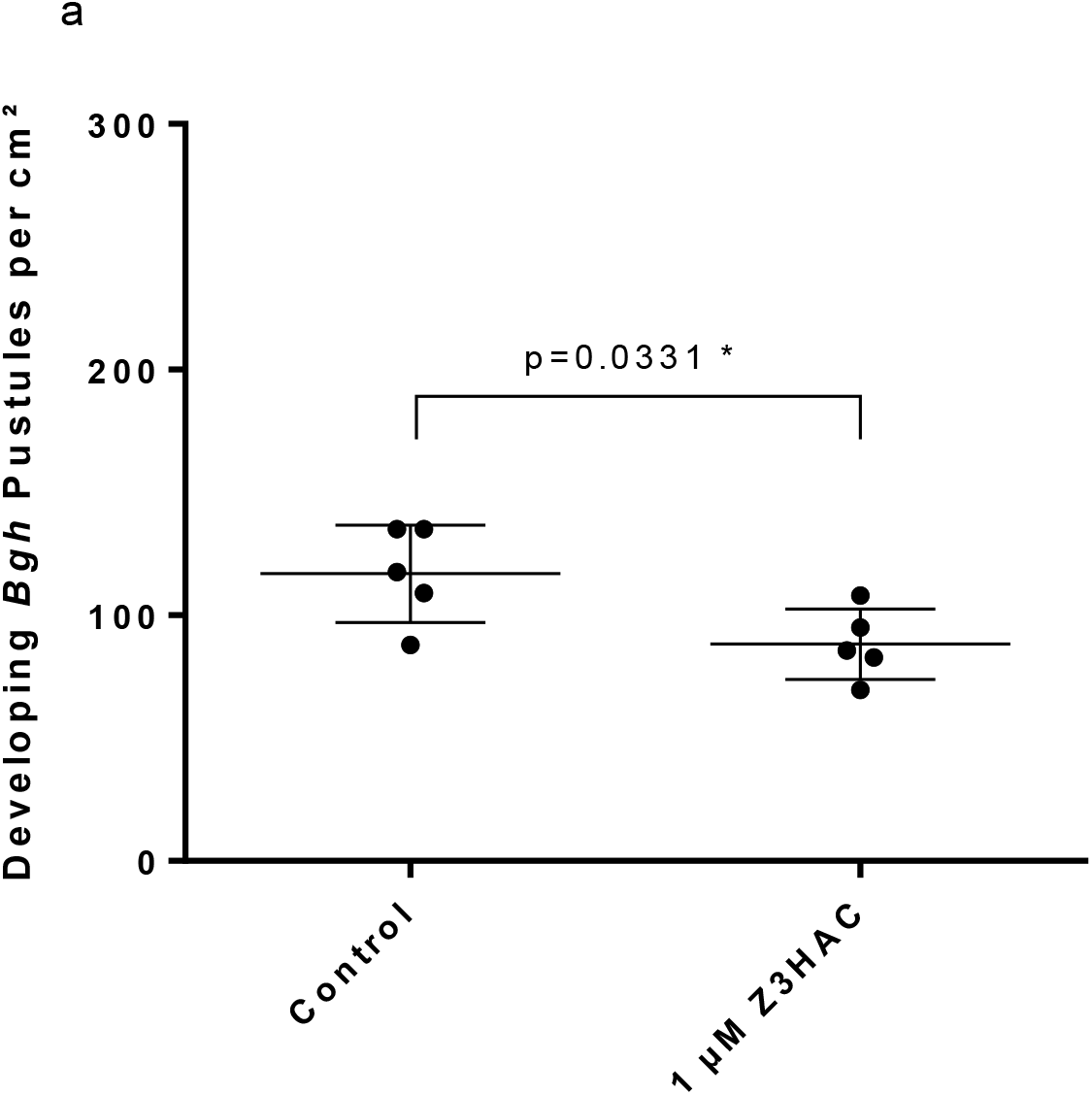
Developing Bgh Pustules on 2^nd^ leaf of young barley plants (cv. Ingrid). After 24 h exposure with 1 µM Z3HAC in a static headspace the individuals were harvested and inoculated with *Bgh* spores on agar plates. 72 h after inoculation microcolonies were counted. Shown are developing *Bgh* pustules per cm^2^ leaf area compared to a non-pre-exposed control. Error bars represent mean ± SD with n=5.

## References

Ameye M., Allmann S., Verwaeren J., Smagghe G., Haesaert G., Schuurink R.C., Audenaert K. (2018) Green leaf volatile production by plants: a meta-analysis. The New phytologist, 220, 666–683.

Ameye M., Audenaert K., Zutter N. de, Steppe K., van Meulebroek L., Vanhaecke L., Vleesschauwer D. de, Haesaert G., Smagghe G. (2015) Priming of wheat with the green leaf volatile Z-3-hexenyl acetate enhances defense against Fusarium graminearum but boosts deoxynivalenol production. Plant physiology, 167, 1671–1684.

Baldwin I.T., Schultz J.C. (1983) Rapid changes in tree chemistry induced by damage: evidence for communication between plants. Science, 227–279.

Bate N., Rothstein S.J. (1998) C6-volatiles derived from the lipoxygenase pathway induce a subset of defense-related genes. The Plant journal, 561–569.

Beßer K., Jarosch B., Langen G., Kogel K.H. (2000) Expression analysis of genes induced in barley after chemical activation reveals distinct disease resistance pathways. Molecular Plant Pathology, 1, 277–286.

Bradford M.M. (1976) A rapid and sensitive method for the quantification of microgram quantities of protein utilizing the principle of protein-dye binding. Analytical Biochemistry, 248–254.

Breiman L. (2001) Random forests. Machine Learing, 5–32.

Delory B.M., Delaplace P., Du Jardin P., Fauconnier M.-L. (2016) Barley (Hordeum distichon L.) roots synthesise volatile aldehydes with a strong age-dependent pattern and release (E)-non-2-enal and (E,Z)-nona-2,6-dienal after mechanical injury. Plant physiology and biochemistry PPB, 104, 134–145.

Dey S., Wenig M., Langen G., Sharma S., Kugler K.G., Knappe C., Hause B., Bichlmeier M., Babaeizad V., Imani J., Janzik I., Stempfl T., Hückelhoven R., Kogel K.-H., Mayer K.F.X., Vlot A.C. (2014) Bacteria-triggered systemic immunity in barley is associated with WRKY and ETHYLENE RESPONSIVE FACTORs but not with salicylic acid. Plant Physiol, 166, 2133–2151.

Dudareva N., Negre F., Nagegowda D.A., Orlova I. (2005) Plant Volatiles. Recent Advances and Future Perspectives. Critical Reviews in Plant Sciences, 417–440.

Eichmann R., Hückelhoven R. (2008) Accommodation of powdery mildew fungi in intact plant cells. Journal of Plant Physiology, 165, 5–18.

Engelberth J., Alborn H., Schmelz E., Tumlinson J. (2004) Airborne signals prime plant against insect herbivore attack. Proceedings of the National Academy of Sciences of the USA, 1781–1785.

Faoro F., Maffi D., Cantu D., Iriti M. (2008) Chemical-induced resistance against powdery mildew in barley: the effects of chitosan and benzothiadiazole. BioControl, 53, 387–401.

Finiti I., La O Leyva M. de, Vicedo B., Gómez-Pastor R., López-Cruz J., García-Agustín P., Real M.D., González-Bosch C. (2014) Hexanoic acid protects tomato plants against Botrytis cinerea by priming defence responses and reducing oxidative stress. Molecular Plant Pathology, 15, 550–562.

Frost C.J., Mescher M.C., Dervinis C., Davis J.M., Carlson J.E., Moraes C.M. de (2008) Priming defense genes and metabolites in hybrid poplar by the green leaf volatile cis-3-hexenyl acetate. The New phytologist, 180, 722–734.

Gfeller A., Laloux M., Barsics F., Kati D.E., Haubruge E., Du Jardin P., Verheggen F.J., Lognay G., Wathelet J.-P., Fauconnier M.-L. (2013) Characterization of volatile organic compounds emitted by barley (Hordeum vulgare L.) roots and their attractiveness to wireworms. Journal of chemical ecology, 39, 1129–1139.

Good, A. G., Crosby W.L. (1989) Induction of Alcohol Dehydrogenase and Lactate Dehydrogenase in Hypoxically Induced Barley. Plant physiology, 90, 860–866.

Harberd N.P., Edwards K.J.R. (1983) Further studies on the alcohol dehydrogenases in barley: evidence for a third alcohol dehydrogenase locus and data on the effect of an alcohol dehydrogenase–1. Genet. Res., 109–116.

Hatanaka A. (1993) The biogeneration of green odour by green leaves. Phytochemistry, 34, 1201–1218.

Hückelhoven R., Kogel K.H. (1998) Tissue-Specific Superoxide Generation at Interaction Sites in Resistant and Susceptible Near-Isogenic Barley Lines Attacked by the Powdery Mildew Fungus (Erysiphe graminis f. sp. hordei). MPMI, 292–300.

IS Compendium (2020) Blumeria graminis (powdery mildew of grasses and cereals). Available from https://www.cabi.org/isc/datasheet/22075.

Kallenbach M., Oh Y., Eilers E.J., Veit D., Baldwin I.T., Schuman M.C. (2014) A robust, simple, high-throughput technique for time-resolved plant volatile analysis in field experiments. The Plant journal for cell and molecular biology, 78, 1060–1072.

Käsbauer C.L., Pathuri I.P., Hensel G., Kumlehn J., Hückelhoven R., Proels R.K. (2018) Barley ADH-1 modulates susceptibility to Bgh and is involved in chitin-induced systemic resistance. Plant physiology and biochemistry PPB, 123, 281–287.

Kennedy R.A., Rumpho M.E., Fox T.C. (1992) Anaerobic Metabolism in Plants. Plant physiology, 100, 1–6.

Kessler A., Baldwin I.T. (2002) Plant responses to insect herbivory: the emerging molecular analysis. Annual review of plant biology, 53, 299–328.

Kishimoto K., Matsui K., Ozawa R., Takabayashi J. (2008) Direct fungicidal activities of C6-aldehydes are important constituents for defense responses in Arabidopsis against Botrytis cinerea. Phytochemistry, 69, 2127–2132.

KIta N., Toyoda H., Shishiyama J. (1981) Chronological analysis of cytological responses in powdery-mildewed barley leaves. Can. J. Bot., 59, 1761–1768.

Kogel K.-H., Langen G. (2005) Induced disease resistance and gene expression in cereals. Cellular Microbiology, 7, 1555–1564.

Kost, Heil (2006) Herbivore-induced plant volatiles induce an indirect defence in neighbouring plants. Journal of Ecology, 94, 619–628.

Kravchuk Z., Vicedo B., Flors V., Camañes G., González-Bosch C., García-Agustín P. (2011) Priming for JA-dependent defenses using hexanoic acid is an effective mechanism to protect Arabidopsis against B. cinerea. Journal of Plant Physiology, 168, 359–366.

Linkmeyer A., Götz M., Hu L., Asam S., Rychlik M., Hausladen H., Hess M., Hückelhoven R. (2013) Assessment and Introduction of Quantitative Resistance to Fusarium Head Blight in Elite Spring Barley. Phytopathology, 1252–1259.

Maffei M.E. (2010) Sites of synthesis, biochemistry and functional role of plant volatiles. South African Journal of Botany, 76, 612–631.

Matsui K. (2006) Green leaf volatiles: hydroperoxide lyase pathway of oxylipin metabolism. Current opinion in plant biology, 9, 274–280.

Mithöfer A., Boland W., Maffei M.E. (2009) Molecular Aspects of Plant Disease Resistance. Chemical Ecology of Plant-Insect-Interaction, 261–291.

Myint T., Ismawanto S., Namasivayam P., Napis S., Abdulla M.P. (2015) Expression analysis of the ADH genes in Arabidopsis plants exposed to PEG-induced water stress. World Journal of Agricultural Research, 57–65.

Nakamura S., Hatanaka A. (2002) Green-Leaf-Derived C6-Aroma Compounds with Potent Antibacterial Action That Act on Both Gram-Negative and Gram-Positive Bacteria. J. Agric. Food Chem, 7639–7644.

Ninkovic V., Markovic D., Rensing M. (2021) Plant volatiles as cues and signals in plant communication. Plant, cell & environment, 44, 1030–1043.

Otto M. (2011) Analytische Chemie, 4.th edn. Wiley-VCH, Germany.

Panstruga R. (2003) Establishing compatibility between plants and obligate biotrophic pathogens. Current Opinion in Plant Biology, 6, 320–326.

Paré P., Tumlinson J. (1996) Plant Volatile Signals in Response to Hervbivore Feeding. Behavioral Ecology Symposium, 93–103.

Pathuri I.P., Reitberger I.E., Hückelhoven R., Proels R.K. (2011) Alcohol dehydrogenase 1 of barley modulates susceptibility to the parasitic fungus Blumeria graminis f.sp. hordei. J Exp Bot, 62, 3449–3457.

Piesik D., Łyszczarz A., Tabaka P., Lamparski R., Bocianowski J., Delaney K.J. (2010) Volatile induction of three cereals: influence of mechanical injury and insect herbivory on injured plants and neighbouring uninjured plants. Annals of Applied Biology, 157, 425–434.

Piesik D., Pańka D., Delaney K.J., Skoczek A., Lamparski R., Weaver D.K. (2011) Cereal crop volatile organic compound induction after mechanical injury, beetle herbivory (Oulema spp.), or fungal infection (Fusarium spp.). Journal of plant physiology, 168, 878–886.

Proels R.K., Westermeier W., Hückelhoven R. (2011) Infection of barley with the parasitic fungus Blumeria graminis f.sp. hordei results in the induction of HvADH1 and HvADH2. Plant Signaling & Behavior, 6, 1584–1587.

Pubchem Open Chemistry Datebase (2016), 8600 Rockville Pike, BethesdaMD, 20894USA.

Scala A., Allmann S., Mirabella R., Haring M.A., Schuurink R.C. (2013) Green leaf volatiles: a plant’s multifunctional weapon against herbivores and pathogens. International journal of molecular sciences, 14, 17781–17811.

Schulze-Lefert P., Vogel J. (2000) Closing the ranks to attack by powdery mildew. Trends in Plant Science, 5, 343–348.

Shiojiri K., Kishimoto K., Ozawa R., Kugimiya S., Arimura G. (2006) Changing green leaf volatile biosynthesis in plants: an approach for improving plant resistance against both herbivores and pathogens. Proceedings of the National Academy of Sciences of the USA, 16672–16676.

Strommer J. (2011) The plant ADH gene family. The Plant journal for cell and molecular biology, 66, 128–142.

Tholl D., Boland W., Hansel A., Loreto F., Röse U.S.R., Schnitzler J.-P. (2006) Practical approaches to plant volatile analysis. The Plant journal for cell and molecular biology, 45, 540–560.

Ton J., D’Alessandro M., Jourdie V., Jakab G., Karlen D., Held M., Mauch-Mani B., Turlings T.C.J. (2007) Priming by airborne signals boosts direct and indirect resistance in maize. The Plant journal for cell and molecular biology, 49, 16–26.

Torres D.P., Proels R.K., Schempp H., Hückelhoven R. (2017) Silencing of RBOHF2 Causes Leaf Age–Dependent Accelerated Senescence, Salicylic Acid Accumulation, and Powdery Mildew Resistance in Barley. Molecular Plant-Microbe Interactions, 906–918.

Trick M., Dennis E.S., Edwards K.J.R., Peacock W.J. (1988) Molecular analysis of the alcohol dehydrogenase gene family of barley. Plant Molecular Biology, 147–160.

Vallelian-Bindschedler L., Schweizer P., Mösinger E., Métraux J.-P. (1998) Heat-induced resistance in barley to powdery mildew (Blumeria graminis f.sp. hordei) is associated with a burst of active oxygen species. Physiological and Molecular Plant Pathology, 185–199.

Vicedo B., Flors V., de la O Leyva, María, Finiti I., Kravchuk Z., Real M.D., García-Agustín P., González-Bosch C. (2009) Hexanoic Acid-Induced Resistance Against Botrytis cinerea in Tomato Plants. MPMI, 1455–1465.

Wiese J., Kranz T., Schubert S. (2004) Induction of pathogen resistance in barley by abiotic stress. Plant biology (Stuttgart, Germany), 6, 529–536.

Wyand R., Brown J.K.M. (2003) Genetic and forma specialis diversity in Blumeria graminis of cereals and its implications for host-pathogen co-evolution. Molecular Plant Pathology, 187.

Xiong L., Schumaker K.S. (2002) Cell Signaling during Cold, Drought, and Salt Stress. The Plant Cell, 14, S165–S183.

Ye M., Liu M., Erb M., Glauser G., Zhang J., Li X., Sun X. (2021) Indole primes defence signalling and increases herbivore resistance in tea plants. Plant Cell Environ, 44, 1165–1177.

